# Repeated phenotypic evolution by different genetic routes: the evolution of colony switching in *Pseudomonas fluorescens* SBW25

**DOI:** 10.1101/414219

**Authors:** Jenna Gallie, Frederic Bertels, Philippe Remigi, Gayle C Ferguson, Sylke Nestmann, Paul B Rainey

## Abstract

Repeated evolution of functionally similar phenotypes is observed throughout the tree of life. The extent to which the underlying genetics are conserved remains an area of considerable interest. Previously, we reported the evolution of colony switching in two independent lineages of *Pseudomonas fluorescens* SBW25 (Beaumont *et al.*, 2009). The phenotypic and genotypic bases of colony switching in the first lineage (Line 1) have been described elsewhere (Beaumont *et al.*, 2009; Gallie *et al.*, 2015). Here, we deconstruct the evolution of colony switching in the second lineage (Line 6). We show that, as for Line 1, Line 6 colony switching results from an increase in the expression of a colanic acid-like polymer (CAP). At the genetic level, nine mutations occur in Line 6. Only one of these - a non-synonymous point mutation in the housekeeping sigma factor *rpoD* - is required for colony switching. In contrast, the genetic basis of colony switching in Line 1 is a mutation in the metabolic gene *carB* (Beaumont *et al*., 2009). A molecular model has recently been proposed whereby the *carB* mutation increases capsulation by redressing the intracellular balance of positive (ribosomes) and negative (RsmAE/CsrA) regulators of a positive feedback loop in capsule expression (Remigi *et al.*, 2018). We show that Line 6 colony switching is consistent with this model; the *rpoD* mutation generates an increase in ribosome expression, and ultimately an increase in CAP expression.

## INTRODUCTION

The repeated appearance of similar phenotypes is a striking feature amid the immense diversity of life. Many phenotypes have evolved multiple independent times in different lineages (Conway Morris, 1999). Examples include the evolution of analogous wing-like structures for flight in pterosaurs, birds, insects and bats (Alexander, 2015), C4 photosynthetic pathways in plants (Sage *et al.*, 2011), and single-lens camera eyes in vertebrates and molluscs (Ogura *et al.*, 2004). An intriguing aspect of repeated phenotypic evolution is the extent to which the underlying genetics are also conserved. It is commonly thought that the degree to which two organisms are related correlates with the degree of genetic parallelism underpinning evolutionary innovations, to the extent that repeated phenotypic evolution has historically been divided into ‘parallel evolution’ (the evolution of one phenotype from similar genetic backgrounds), and ‘convergent’ evolution (the evolution of one phenotype from distantly related organisms). The accumulation of genetic data in recent years has shown this assumption to be in need of revision. For example, clonal populations of *Escherichia coli* adapt to thermal stress *via* different genetic routes (Riehle *et al.*, 2001), while pigmentation changes in mice and lizards are both underpinned by mutations in the *Mc1r* gene (Nachman *et al.*, 2003; Rosenblum *et al.*, 2004). The increasing number of examples of disparity between degree of relatedness and genetic parallelism (reviewed in Arendt and Reznick, 2008) hints at the underappreciated and poorly understood complexity of biological systems.

An evolution experiment with populations of the model bacterium *Pseudomonas fluorescens* SBW25 (Beaumont *et al.*, 2009) has provided the opportunity to characterize a case of repeated phenotypic evolution in unusual detail. Twelve independent populations were subjected to multiple rounds of selection for novel colony morphologies. In every population, each round of selection concluded with the isolation of a single colony that had a phenotype different to that of the immediate ancestor for continuation into the subsequent round of selection (**fig. 1A**). The final result was 12 independent evolutionary lineages, each with a clearly defined history of colony phenotypes and underlying genetic changes. Two lineages (Lines 1 and 6) converged on a similar striking capacity to stochastically switch – at high frequency – between different colony morphologies.

**Fig. 1.**
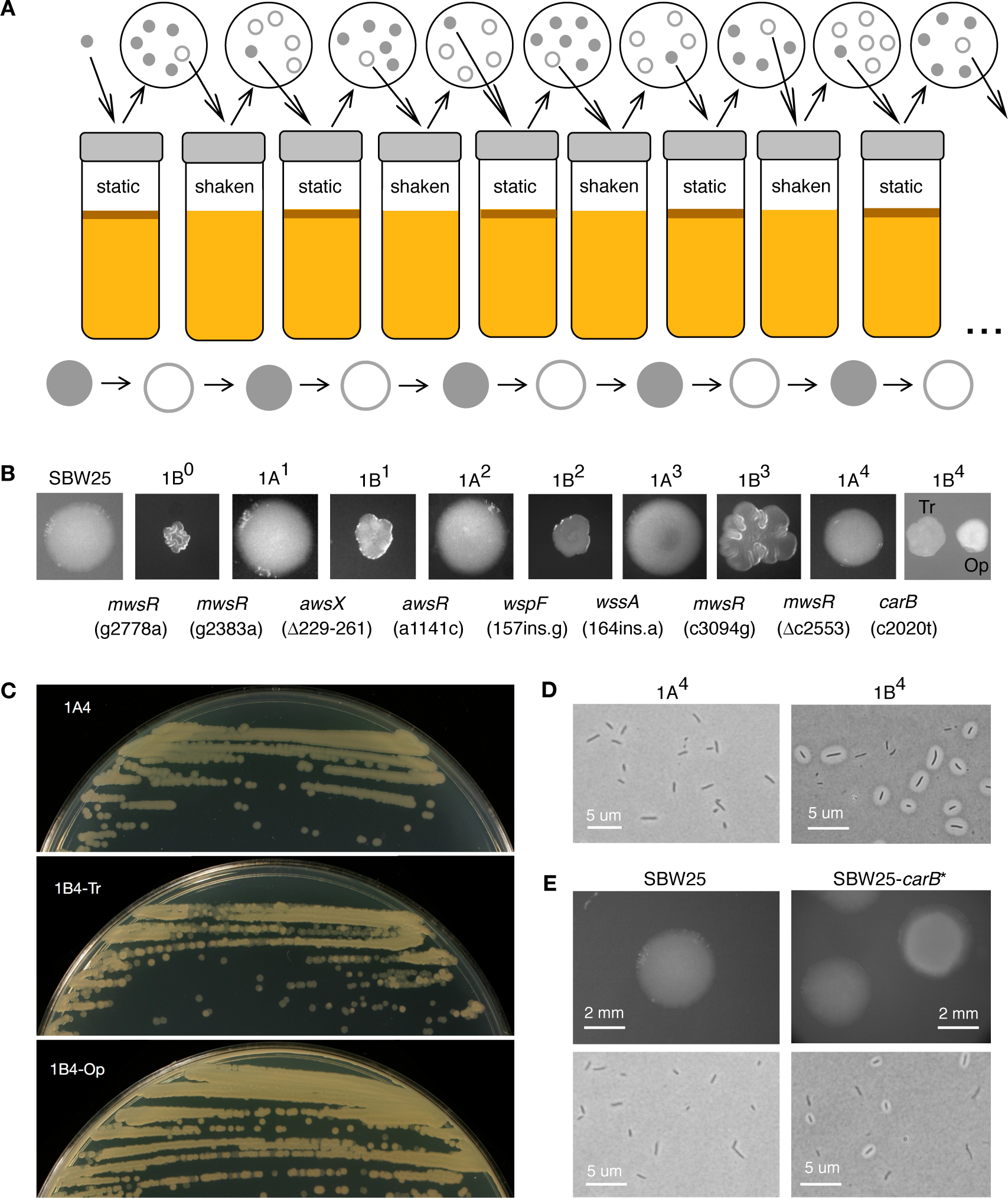
The reverse evolution experiment (REE) and the emergence of colony switching in Line 1. **(A)** Cartoon of a single line of the 12-line REE (Beaumont *et al*., 2009). Populations were subjected to bouts of selection in static or shaken liquid KB. After each bout cells were plated on KB agar and a single colony with novel morphology was used to start the subsequent round in the opposite environment. **(B)** Each of the nine strains in the Line 1 evolutionary series (SBW25→1B^4^) has a colony phenotype distinct from that of its immediate ancestor. New mutations are noted as “*gene* (mutation)” at the point of occurrence. Tr=translucent, Op=opaque. **(C)** Bi-directional colony switching in 1B^4^. 1A^4^ colony streaked onto a fresh plate (top) generates colonies of a single type, while a 1B^4^ Tr (middle) or Op colony (bottom) generates a mixture of colony types. **(D)** 1B^4^ cells are either capsulated (Cap^+^) or non-capsulated (Cap^-^), while 1A^4^ cells are generally Cap^-^. **(E)** SBW25 colonies are uniform while SBW25-*carB*\* (SBW25 into which the c2020t *carB* mutation is engineered) shows colony bistability. All colonies grown on KB agar (26^°^C, 48 hrs); cells grown in shaken KB for 16-24 hrs before staining with India ink. Exposure and brightness of some images altered in Preview.

Colony switching in Line 1 has been extensively investigated (Beaumont *et al.*, 2009; Rainey *et al.*, 2011; Libby and Rainey, 2011; Gallie *et al.*, 2015; Remigi *et al.*, 2018). Emergent genotype 1B^4^ produces a mixture of opaque and translucent colonies, and a corresponding mixture of capsulated and non-capsulated cells (**fig. 1B**, **1C;** Beaumont *et al.*, 2009). The capsule consists of a colanic acid-like polymer (CAP), the ON/OFF expression of which leads to colony switching (Gallie *et al.*, 2015). Nine mutational steps occurred during the evolution of 1B^4^ (**fig. 1B**). The first eight occur in genes involved in the production of c-di-GMP, a secondary messenger that affects the expression of a well-characterized acetylated cellulosic polymer (ACP, cellulose; Spiers *et al.*, 2003; McDonald *et al.*, 2009). The final mutation affected the central metabolic gene *carB* (c2020t; amino acid change R674C). This mutation, which is alone sufficient to cause colony switching, perturbs intracellular pyrimidine pools (Gallie *et al.*, 2015; **fig. 1D)**. Pyrimidine deficiency in 1B^4^ has recently been shown to generate an increase in intracellular ribosome concentration, leading to the proposal of a translational control model for capsule switching (Remigi *et al.*, 2018; see also **fig. 6A**). Briefly, the model proposes that capsule switching results from competition for binding sites on the mRNA of *pflu3655-pflu3657*, which encodes transcriptional regulators of CAP biosynthetic genes; ribosome binding results in translation (and promotion of capsulation), while RsmAE/CsrA binding inhibits translation (favouring the non-capsulated state; Remigi *et al.*, 2018). The ribosome increase in 1B^4^ is expected to tip the balance of the switch in favour of translation, increasing the probability of capsulation.

In this work, we characterize the phenotypic and genetic bases of colony switching in the second emergent genotype, 6B^4^. Comparisons with 1B^4^ demonstrate that 6B^4^ colony switching is a very similar phenotype realized by a different genetic route. We also show that the two genetic routes are reconciled at the molecular mechanistic level.

## RESULTS

### 6B^4^ shows colony and capsule instability

The evolutionary history of 6B^4^ includes ten colony phenotypes, with translucent-opaque colony instability emerging after nine rounds of evolution (**fig. 2A**). 6B^4^ colonies are comprised of a mixture of capsulated and non-capsulated cells with 6B^4^ populations containing a significantly higher proportion of capsulated cells than its immediate ancestor 6A^4^ (**fig. 2B**, **2C**; Welch two-sample *t*-test *p*=1.3×10^-4^). Single 6B^4^ cells of either type give rise to mixed Cap^+^/Cap^-^ populations (**supplementary text S1**). We conclude that colony switching in 6B^4^ has the same underlying phenotypic basis as in 1B^4^: the ON/OFF switching of capsule biosynthesis. However, under the conditions tested, the proportion of capsulated cells is significantly higher in 6B^4^ than 1B^4^ populations (**fig. 2C**; Welch two-sample *t*-test 1.9×10^-4^). Additionally, on KB agar 6B^4^ capsules are between 1.26 and 1.83 times larger than those in 1B^4^ (*t*-test for no difference in capsule area *p*=9.602×10^-10^), despite no difference in cell size between the two strains (*t*-test for no difference in cell area *p*=0.5236; see **supplementary text S1**).

**Fig. 2.**
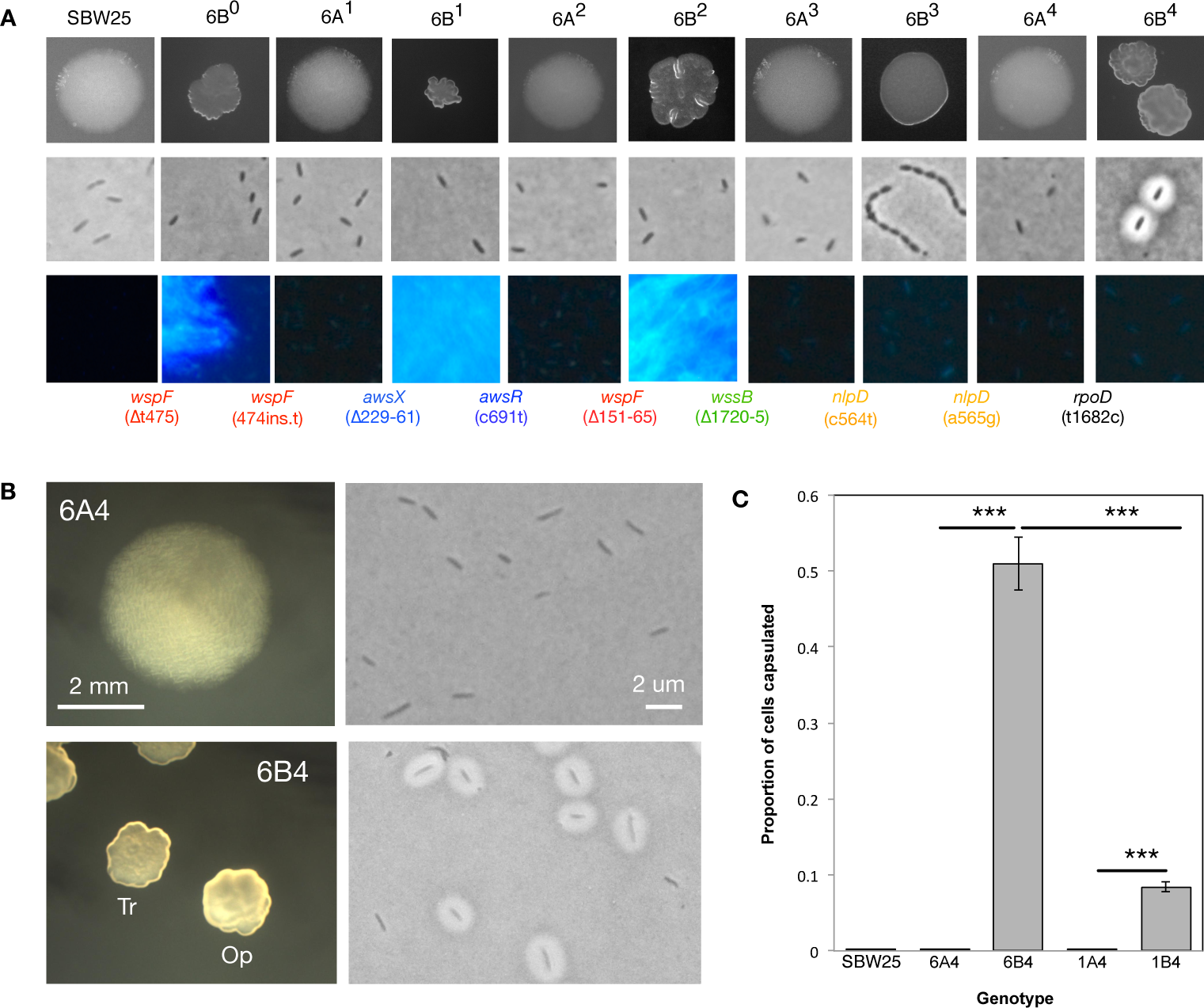
Emergent capsule switching in Line 6 of the REE. **(A)** The phenotypes and genotypes of the Line 6 evolutionary series: colony morphology on KB agar (first row), morphology of cells grown in shaken KB microcosms and stained with India ink (second row), ability to produce ACP (cells grown on KB agar with 5% calcofluor; third row), and new mutations acquired (mutations are shown at the point of occurrence by listing the gene and the mutation underneath in parentheses. Genes belonging to the same locus share a font colour; fourth row). **(B)** Colony and cell morphologies of 6B^4^ and its non-switching immediate ancestor, 6A^4^. 6B^4^ gives rise to translucent (Tr) and opaque (Op) colonies, plus capsulated and non-capsulated cells. **(C)** The proportion of capsulated cells in populations of various genotypes. Each bar represents the mean of five replicate populations grown overnight in KB microcosms. Error bars represent one standard error and stars denote statistical significance (* * * =*p*<0.001). Brightness, contrast and/or saturation of some images altered in Preview.

### 6B^4^ capsule expression is due to transcriptional regulation of wcaJ-wzc

In order to identify the genetic basis of the 6B^4^ capsule, 6B^4^ was subjected to transposon mutagenesis. In a screen of ∼10,000 transposon mutants, 55 with altered levels of capsulation were identified, and the transposon insertion site determined for each (**supplementary table S1**). Microscopic screening of cells showed capsule production to be eliminated in 43 genotypes, and severely reduced in a further nine genotypes. Three genotypes showed an increase in capsule production.

Of the genotypes with eliminated or reduced capsule production, 41 (75%) contained insertions in genes required for the production of CAP, a polymer previously described as the structural basis of the 1B^4^ capsule (Gallie *et al.*, 2015). These include insertions in genes predicted to encode CAP precursor biosynthetic machinery (*e.g., algC*), CAP biosynthetic machinery (20 genes: *wcaJ-wzc*) and probable CAP regulators (*pflu3655, pflu3656, pflu3657, gacA/gacS*). A direct deletion of the CAP biosynthetic locus from 6B^4^ resulted in loss of both cell capsulation and colony instability (**fig. 3A**, **supplementary text S2**). Together these results demonstrate that the structural basis of the 6B^4^ capsule is encoded by the *wcaJ-wzb* locus.

**Fig. 3.**
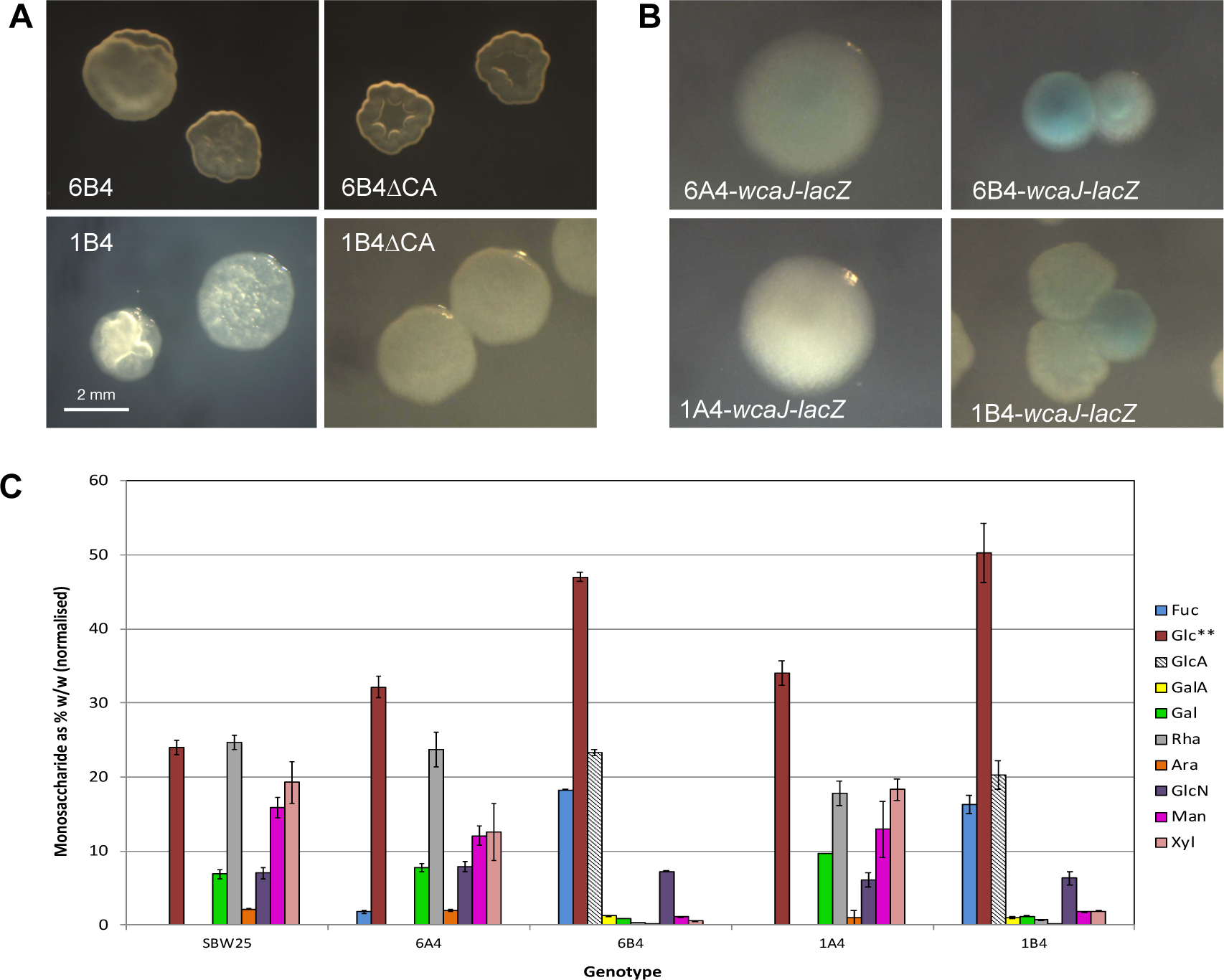
The structural basis of the 6B^4^ capsule is a colanic acid-like polymer (CAP). (**A**) Deletion of *wcaJ-wzb* (the ∼23.3 kb CAP-biosynthetic locus) results in loss of capsule switching. (**B**) Colonies in which a transcriptional fusion of *lacZ* has been made to CAP biosynthetic gene *wcaJ* in the 6A^4^, 6B^4^, 1A^4^ and 1B^4^ backgrounds (on LB+Tc+X-gal plates, 42 hours). 6B^4^ and 1B^4^ develop a mixture of blue and white colonies, showing a degree of transcriptional control of *wcaJ*. Exposure of some images altered in Preview. (**C**) An analysis of the monosaccharide components of the extracellular polysaccharide produced by 6B^4^, 1B^4^ and ancestors shows the emergent production of a similar polymer in both 6B^4^ and 1B^4^ (see also **supplementary table S2**). Bars are the mean of two replicate measurements, and error bars are 1 standard error. The components of this polymer are consistent with a colanic acid-like polymer (CAP). There are also two unidentified sugar components found in 6B^4^ and 1B^4^ (not shown).

A transcriptional fusion of *lacZ* to *wcaJ* was constructed in 6B^4^ and the phenotype analysed on LB+X-gal plates (**fig. 3B**, **supplementary text S2**). The resulting 6B^4^-*wcaJ*-*lacZ* colonies were a mixture of white (with a high proportion of Cap^-^ cells), and blue (with a high proportion of Cap^+^ cells). The same construction in 6A^4^ – the immediate ancestor – resulted in uniform colonies, showing that CAP expression is at least partially controlled at the level of transcription (later corroborated by RNA-seq data; see **supplementary tables S3-S5**).

### The structural basis of the 6B^4^ capsule is CAP

In order to directly investigate the composition of the 6B^4^ capsule, extracellular polysaccharide (EPS) was extracted from SBW25, 6A^4^, 6B^4^, 1A^4^ and 1B^4^, and the component sugars from each strain analysed by chromatography (results for SBW25, 1A^4^ and 1B^4^ reported previously in (Gallie *et al.*, 2015; **fig. 3C**, **supplementary table S2**). The analysis shows differences in the expression of several components: in 6B^4^ relative to 6A^4^, the expression of D-Fucose (Fuc), D-glucuronic acid (GlcA), D-Galactouronic acid (GalA) and two unknowns are increased. Each of these is also increased in 1B^4^ relative to 1A^4^, indicating that the 1B^4^ and 6B^4^ capsule polymers are the same.

Thus far, the transposon mutagenesis, strain constructions and structural analysis of the capsule polymers (and later, RNA-seq data) point to the same phenotype for 1B^4^ and 6B^4^: switching between opaque and translucent colonies caused, at the single cell level, by ON/OFF expression of CAP. The only difference observed between the two genotypes lies in the frequency of capsulation and size of capsules (both increased in 6B^4^ relative to 1B^4^).

### Nine mutations detected in 6B^4^

Next, the genetic basis of 6B^4^ capsule switching was investigated. Whole genome sequencing of 6B^4^ identified seven mutations. This was surprising, as nine mutations were expected – one *per* round of REE selection (see **fig. 1A**). Sanger sequencing of the identified loci at each bottleneck in the evolutionary series revealed two gaps at the beginning: SBW25→6B^0^ (selection round 1) and 6B^0^→6A^1^ (selection round 2; **table 1**, **fig. 2A**). Extensive previous knowledge suggested that these two genotypes almost certainly carried mutations in one of three loci (*wsp, aws, mwsR*) (McDonald *et al.*, 2009). Sanger sequencing of *wspF* revealed a point deletion in 6B^0^ (**Δ**t475) that was absent in 6A^1^ (**fig. 2A**). The loss of the *wspF* **Δ**t475 mutation was not repeated among twenty independent repeats of a single round of REE from 6B^0^. The absence of the *wspF* deletion in 6A^1^ most likely represents a rare mutational reversal (**table 1**, **fig. 2A**).

**Table 1:**
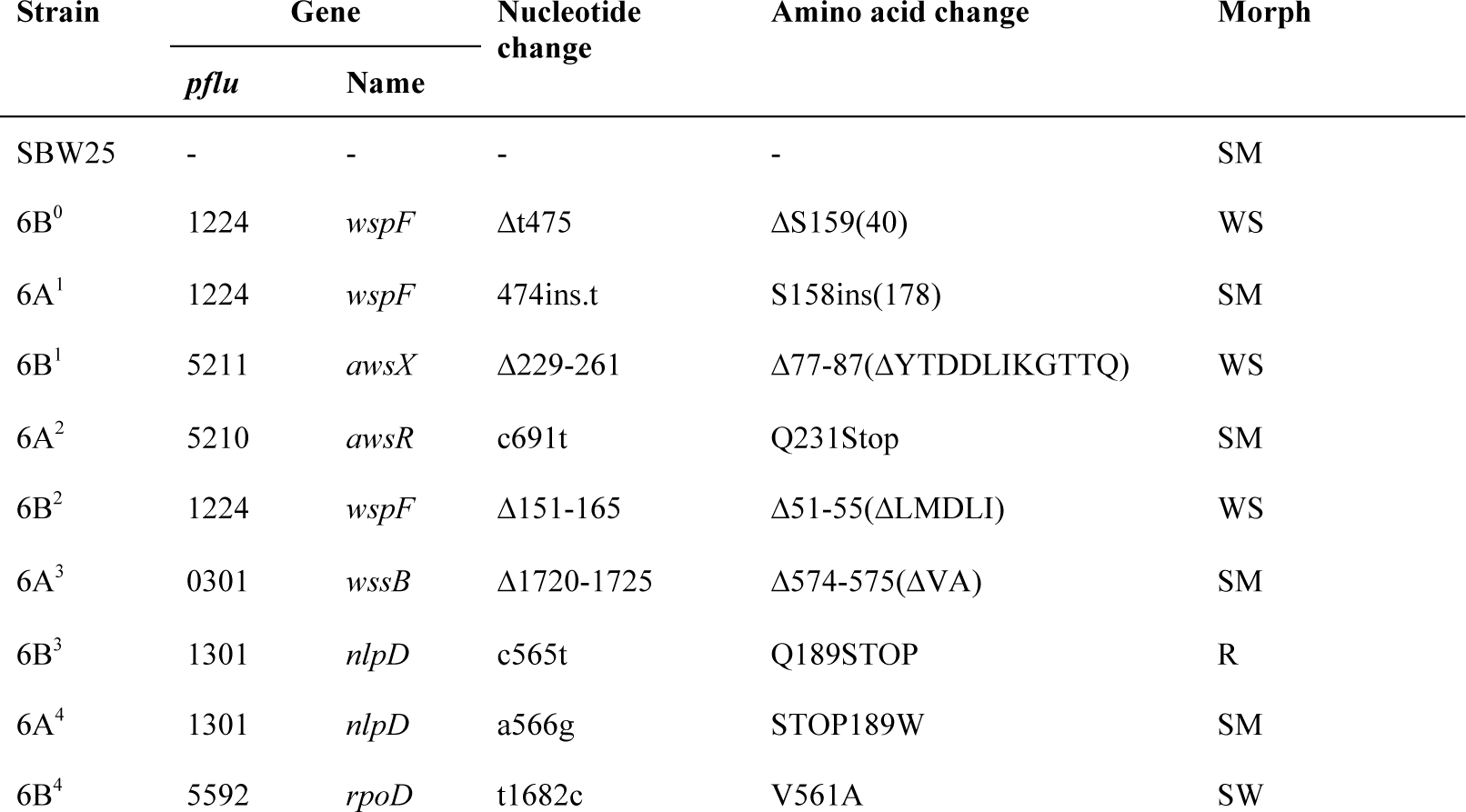
Details and chronological order of mutations in the Line 6 evolutionary series.

The nine Line 6 mutations occur in a modular, paired fashion. The first six mutations occur in previously identified c-di-GMP producing loci (*awsX/awsR, wspF/wssB*); mutations in these loci are known to cause the gain and loss of cellulose production and wrinkly spreader colony morphology (McDonald *et al.*, 2009; Beaumont *et al.*, 2009; Gallie *et al.*, 2015; Lind *et al.*, 2015; Lind *et al.*, 2016). The sixth mutation – an in-frame, six base pair deletion in the cellulose biosynthetic gene *wssB* – completely abolishes cellulose production (**fig. 2A**), and appears to render downstream genotypes unable to produce the wrinkly spreader colony morphology by further mutation. Accordingly, the next pair of mutations occur in an unrelated locus: *nlpD* (*pflu1301*), which encodes a lipoprotein predicted to have a function in cell wall formation and cell separation in a range of bacteria (Stohl *et al.*, 2015; Lind *et al.*, 2016; Tsang *et al.*, 2017; Yang *et al.*, 2017). The first of these, mutation seven, generates a nonsense mutation in *nlpD* resulting in the production of cell chains and round colonies in 6B^3^ (**fig. 2A**). This mutation has previously been reported to generate a cell chain phenotype in SBW25 (Lind *et al.*, 2016), and similar mutations have been reported in *Escherichia coli* (Uehara *et al.*, 2010), *Vibrio cholerae* (Möll *et al.*, 2014) and *Yersinia pestis* (Tidhar *et al.*, 2009). In short, NlpD is an activator of cell division protein AmiC; inactivation of NlpD leads to incomplete cell division. Mutation eight converts the *nlpD* nonsense mutation into a tryptophan residue, reversing the cellular and colony phenotypes in 6A^4^ (**fig. 2A**). The final mutation, with which colony switching emerges, is in *rpoD* (t1682c, resulting in amino acid change V561A). This gene encodes the housekeeping sigma factor (Σ^70^) that controls transcription of many genes involved in cell growth and division (Schulz *et al.*, 2015).

There are two notable points of similarity and contrast between the evolutionary histories of 6B^4^ (**fig. 2A**) and 1B^4^ (**fig. 1B**). Firstly, both lineages begin in a similar fashion with mutations affecting cellulose production and wrinkly spreader colony morphology. In Line 6, mutational routes to the wrinkly spreader phenotype are presumably rendered inaccessible by the sixth mutation (in *wssB*), providing an opening for a pair of mutations in *nlpD*. Contrastingly, cellulose production is not abolished in Line 1, with 1B^4^ staining positive for cellulose and forming wrinkly spreader-like mats. Accordingly, Line 1 mutations are in cellulose-affecting loci up until the final, switch-causing mutation. Secondly, the final mutation in each lineage – that with which colony switching emerges – is a non-synonymous point mutation in different and, at first glance, functionally unrelated housekeeping genes.

### The rpoD t1682c mutation alone generates some capsulation

To confirm that the final mutation causes colony switching in the presence of the other eight mutations, the t1682c *rpoD* mutation was engineered into the immediate ancestor (6A^4^). The resulting genotype, 6A^4^-*rpoD\**, gave rise to high-level colony and CAP switching, showing the same proportion of capsulated cells as 6B^4^ (*t*-test *p*=0.45; **fig. 4**). The *rpoD* mutation was then engineered into the distant ancestor, SBW25, in the absence of any other mutations. The resulting genotype, SBW25-*rpoD*\*, also showed distinct colony types and a high level of capsulation. A capsule counting assay revealed that while the *rpoD* mutation alone was sufficient to cause switching, SBW25-*rpoD*\* showed a lower degree of capsulation than 6B^4^ (*t*-test *p*=1.6×10^-3^; **fig. 4C**). Therefore, while the *rpoD* mutation does cause CAP switching, one or more of the other six mutations contribute quantitatively to the capsule switching phenotype. This is in contrast to the c2020t *carB* mutation in Line 1, which accounts alone for the level of CAP switching seen in 1B^4^ (Gallie *et al.*, 2015).

**Fig. 4.**
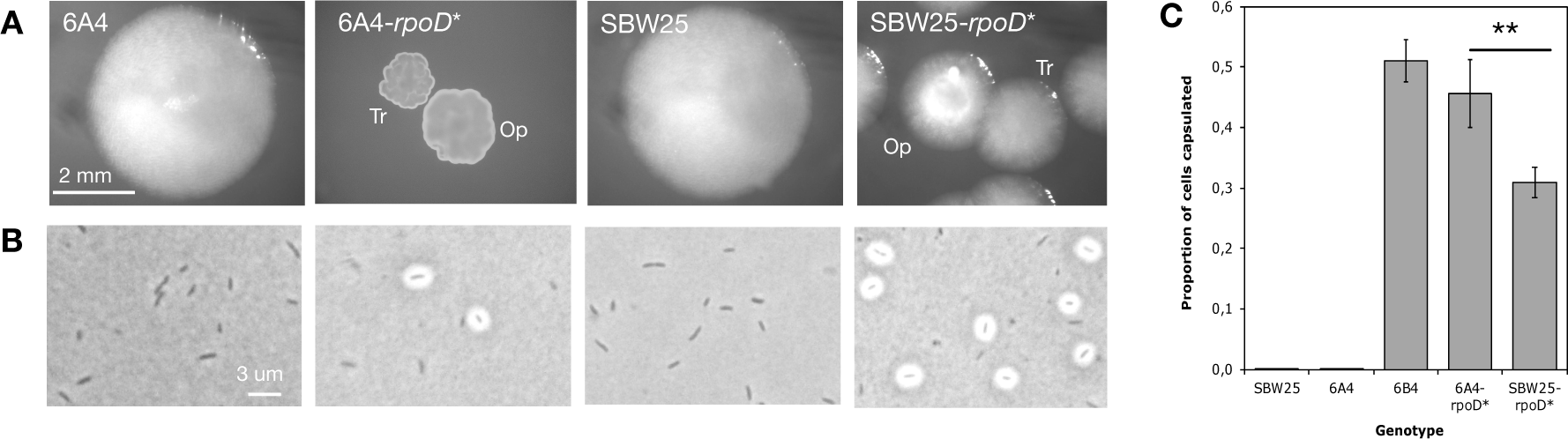
The t1682c *rpoD* mutation causes the emergence of capsule switching both in the presence and absence of the other Line 6 mutations. (**A**) Colony phenotypes on KB agar after 48 hours show that engineered strains carrying the *rpoD* mutation develop colonies of mixed Tr and Op phenotypes. (**B**) Cell phenotype of ancestral non-switching genotypes (6A^4^, SBW25) versus engineered switcher genotypes (6A^4^-*rpoD*\*, SBW25-*rpoD*\*) when grown overnight in KB glass microcosms. Saturation and brightness of photographs altered in Preview. (**C**) The proportion of capsulated cells in populations of various genotypes. Each bar represents the mean of five replicate populations grown overnight in KB glass microcosms. Error bars represent one standard error and stars show statistical significance (* * =0.01<*p*<0.001).

### Repeated evolution of switcher genotypes reveals additional rpoD mutations

To identify additional mutations able to cause capsule switching in 6A^4^, new switcher genotypes were evolved from 6A^4^. Each of 56 independent microcosms was inoculated with 6A^4^ and put through a single round of the REE (Beaumont *et al.*, 2009). Nine new switcher genotypes were isolated from nine independent microcosms (genotypes Re1-Re9; see **supplementary text S1**). Sequencing of *rpoD* revealed a single, non-synonymous point mutation in each; eight of the new switchers (Re1-Re8) contained mutation a1723c leading to amino acid change T575P, while one (Re9) carried a1745c causing amino acid change Q582P. All three *rpoD* mutations (t1682c, a1723c, a1745c) are located in the H-T-H motif that interacts with the -35 consensus sequence of Σ^70^ dependent promoters (**fig. 5A**; Hu and Gross, 1988; Siegele *et al.*, 1989). Interestingly one mutation, T575P, leads to a significantly higher capsulation rate (*t*-test *p*<0.001; **fig. 5B-C**).

**Fig. 5.**
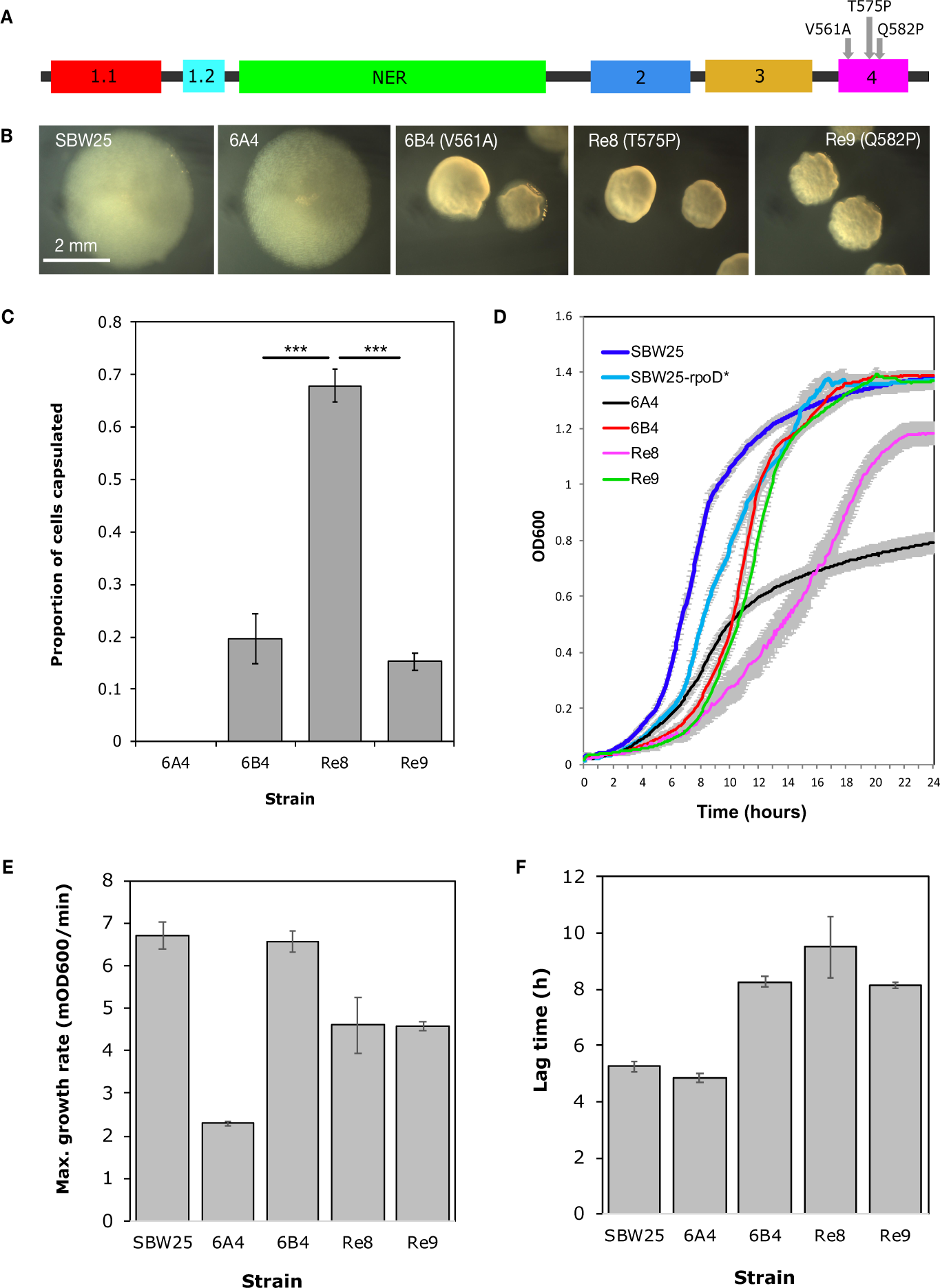
Three *rpoD* mutations have different effects on capsulation and growth. Three amino acid substitutions leading to switching have been identified in Σ^70^. Each of these changes occurs in region 4, which encodes a H-T-H motif that binds to the -35 consensus sequence of Σ^70^-dependent promoters. NER=non-essential region (**A**). In each case, the amino acid substitution leads to the emergence of two colony phenotypes or sectored colonies (**B**; colonies grown on KB agar for ∼56 hours) and an increase in capsulation during a capsule counting assay (**C**; bars are the mean of five replicates). (**D**) 24-hour growth curves in shaken KB medium at 28^°^C. Measurements were taken every 5 minutes, with eight replicates for each strain (against KB blanks). Mean maximum growth rates (**E**) and lag times (**F**) were calculated using a sliding window of six data points. Error bars on all graphs show one standard error.

Changes in such a central gene are expected to have major effects on cell growth. Indeed, the growth profiles of each *rpoD* mutant differ from those of the ancestral strains (**fig. 5D**). The *rpoD* mutations increase growth rate and final density – at the cost of a longer lag phase – in comparison to 6A^4^ (**fig. 5D-F**). These results are consistent with the *rpoD* mutations affecting Σ^70^ activity during exponential growth.

### Gene expression differences in the presence of the t1682c rpoD mutation (RNA-seq)

Changes in the Σ^70^-promoter recognition and binding domain are expected to affect expression from Σ^70^-dependent promoters (or a subset thereof). Thus the effect of the t1682c *rpoD* mutation on intracellular mRNA pools was investigated. Total mRNA was isolated from three biological replicates of each of 6A^4^, 6B^4^-Cap^-^ and 6B^4^-Cap^+^. RNA-seq was performed and three comparative analyses generated: (A) 6A^4^ versus 6B^4^-Cap^-^, (B) 6A^4^ versus 6B^4^-Cap^+^, and (C) 6B^4^-Cap^-^ versus 6B^4^-Cap^+^. A list of genes with detectably different expression levels (∼98% of all predicted genes in the SBW25 genome (Silby *et al.*, 2009)) was generated for each comparison, and the three lists were then further split into genes with and without statistically significantly different expression levels (**supplementary tables S3-S5**).

The greatest number of genes showing statistically significantly different expression was found in comparison B, 6A^4^ versus 6B^4^-Cap^+^, indicating that these are the two most physiologically distinct morphotypes. Of the 1,438 genes identified, 612 were more highly expressed in the ancestral 6A^4^ (including 33 flagella biosynthetic genes), and 826 were more highly expressed in 6B^4^-Cap^+^ (including 24 CAP and seven alginate biosynthetic genes). Comparison A, 6A^4^ versus 6B^4^-Cap^-^, identified 495 differentially expressed genes with statistical significance, 427 (86 %) of which are shared with comparison B. Comparison of 6B^4^-Cap^-^ and 6B^4^-Cap^+^ identified 82 significantly differently expressed genes, 52 of which are more highly expressed in 6B^4^-Cap^-^ (including 30 flagella biosynthetic genes) and 30 in 6B^4^-Cap^+^ (including 12 CAP genes). Notably, mutant *rpoD* was found to be ∼1.74 times more highly expressed in 6B^4^-Cap^+^ than the wild type *rpoD* counterpart in 6A^4^ (*p*=0.0324) indicating that the t1682c *rpoD* mutation leads to activation of *rpoD* transcription and/or inhibition of mRNA degradation (**supplementary table S4**). Levels of *rpoD* mRNA in 6B^4^-Cap^-^ are intermediate between those in 6A^4^ and 6B^4^-Cap^+^, as no significant difference in *rpoD* mRNA levels was detected between 6B^4^-Cap^-^ and either of the other two types (**supplementary tables S3** and **S5**). A further five putative sigma factors are more highly expressed in 6B^4^-Cap^+^ than 6A^4^ (*rspL, pflu2609, pflu2725, pflu3898, pflu4613*), indicating a general shift in the intracellular gene expression between 6A^4^ and 6B^4^-Cap^+^.

The equivalents of the above comparisons have been previously published for Line 1 (GEO database submission number GSE48900; (Gallie *et al.*, 2015). While the numbers of differentially expressed genes are much higher in the Line 1 comparisons – most likely attributable to there being only a single biological replicate for each Line 1 morphotype – the overall pattern remains; the highest number of differentially expressed genes is between 1A^4^ and 1B^4^-Cap^+^, and the lowest between 1B^4^-Cap^-^ and 1B^4^-Cap^+^. A “comparison of comparisons” was performed, whereby each of comparisons A, B and C for Line 6 was equated to the Line 1 counterpart. Lists of shared and unique genes for comparisons A, B and C were generated (**supplementary table S6**). For comparison C, B^4^-Cap^-^ versus B^4^-Cap^+^, 26 genes are common between Line 6 and Line 1; nine of these are more highly expressed in Cap^-^ forms compared to the Cap^+^, and include four flagella genes and five genes of unknown function. The remaining 17 genes are more highly expressed in the Cap^+^ forms than the Cap^-^, and include seven CAP genes, a transcriptional regulator, an inorganic ion transport gene and eight genes of unknown function. Together, the results corroborate the finding that capsules and flagella are mutually exclusive. A similar finding has recently been reported in *Cronobacter sakazakii*, in which induction of colanic acid biosynthesis is accompanied by a reduction in flagella gene expression (Chen *et al.*, 2018).

### Ribosomal genes and proteins are overexpressed in 6B^4^

The recently proposed ribosome-RsmAE model of 1B^4^ capsule switching postulates that capsulation is controlled by the combined intracellular pool of ribosomes and RsmAE (Remigi *et al.*, 2018); **fig. 6A**). According to the model, ribosomes and RsmAE compete for binding of the RBS in *pflu3655-pflu3657* mRNA (which encodes transcriptional activators of the capsule biosynthetic genes); ribosome binding results in translation, activation of a positive feedback loop, and capsulation. RsmAE binding results in inhibition of translation and the non-capsulated state. The model predicts that genotypes with increased capsulation (such as 6B^4^) contain higher levels of *pflu3655-3657* mRNA as a result of a net increase in ribosomes.

**Fig. 6.**
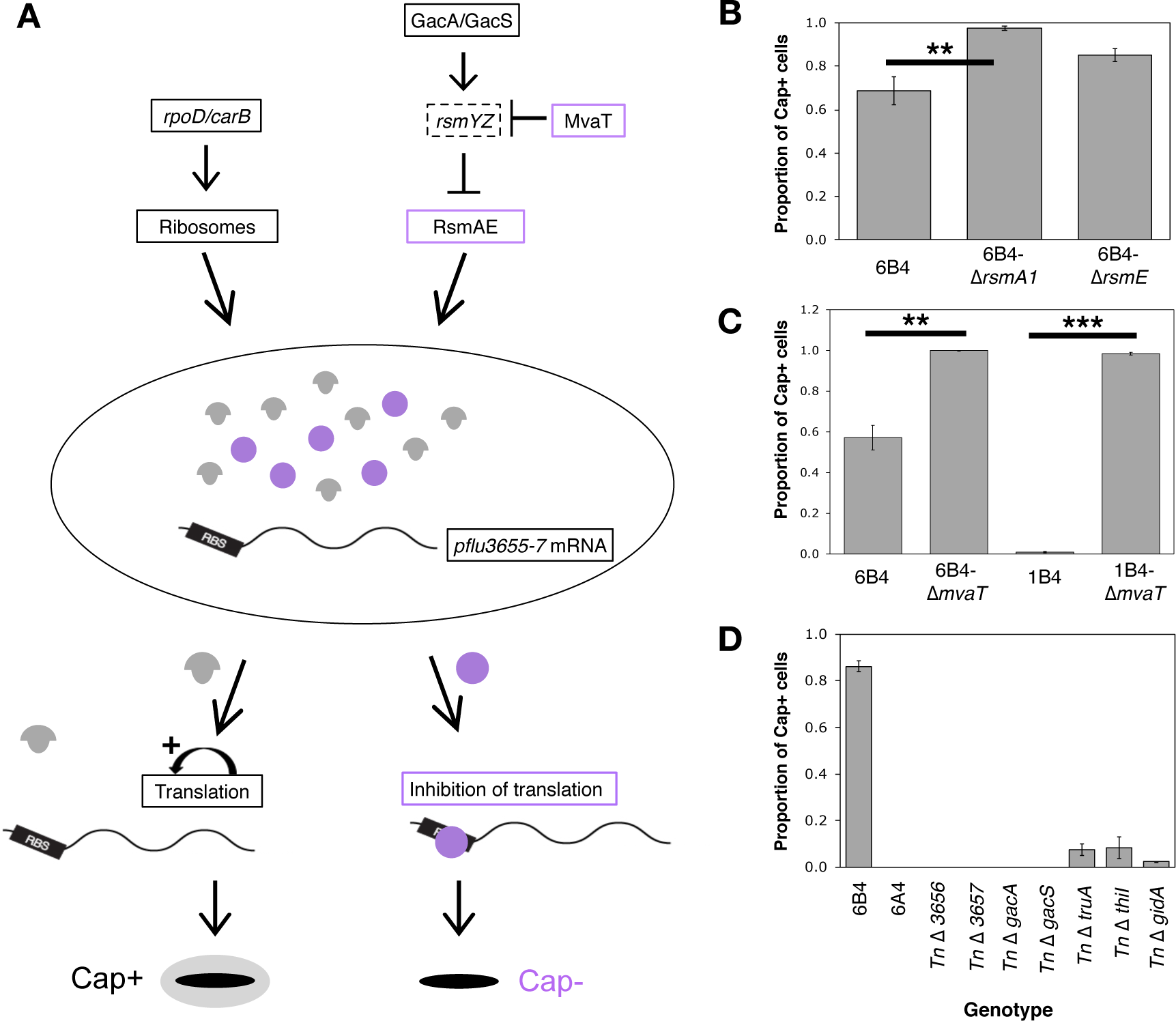
Testing the ribosome-RsmAE model of capsule switching in 6B^4^. **(A)** The model postulates that capsulation is controlled by the combined intracellular pool of ribosomes (grey mushrooms) and RsmAE (purple circles), which compete for binding of *pflu3656-3657* mRNA (encoding transcriptional activators of CAP); ribosome binding results in translation, activation of a positive feedback loop, and capsulation. RsmAE binding results in inhibition of translation and the non-capsulated state (Remigi *et al*., 2018). Intracellular components predicted to have positive or negative influences on capsulation are outlined in black or purple, respectively. Solid outlines indicate components for which evidence has been obtained (transposon mutagenesis, genetic engineering and/or mRNA-seq), dotted lines are used for components for which no evidence has been obtained. The model poses *rpoD* and *carB* as activators of ribosome production. (**B**) Deletion of *rsmA1* from 6B^4^ leads to a significant increase in capsulation (*p*=1.112×10^-3^). (**C**) Deletion of *mvaT* (*pflu4939*) from 6B^4^ or 1B^4^ leads to a significant increase in capsulation (Wilcoxon rank sum test *p*=0.009761; two sample *t*-test *p=*1.952×10^-15^, respectively). (**D**) A capsule counting assay with non-polar, transposon-derived insertions in *pflu3656, pflu3657*, negative regulators of RsmAE (*gacA, gacS*) and cell-wide translation (*truA, gidA, thiI*) lead to a reduction in 6B^4^ capsulation. Bars are the mean of five (B, C) or three (D) replicates, and error bars represent 1 standard error.

Consistent with the model, *pflu3655, pflu3656* and *pflu3657* mRNA levels are significantly higher in 6B^4^-Cap^+^ and 6B^4^-Cap^-^ than in 6A^4^. Indeed *pflu3655* is the most highly differentially expressed gene in all three RNA-seq comparisons: it is expressed 572 fold more highly in 6B^4^-Cap^+^ than 6A^4^, 31 fold in 6B^4^-Cap^-^ versus 6A^4^, and 19 fold in 6B^4^-Cap^+^ versus 6B^4^-Cap^-^ (**supplementary tables S3-S5**). Furthermore the *rpoD* mutation leads to an increase in the mRNA level from ribosomal genes (**fig. 6A**); 40 and 43 (of 53) ribosomal protein genes show increased expression in 6B^4^-Cap^+^ compared to 6A^4^ and 6B^4^-Cap^-^, respectively (**supplementary table S7**). While only three of these show statistical significance (*rpmG, rpmB* and *rpsT* are more highly expressed in 6B^4^-Cap^+^ than 6A^4^; **supplementary text S3**), there is a clear pattern of higher expression in the capsulated form. Further binomial tests provide strong evidence for increased expression of ribosomal genes in 6B^4^-Cap^+^ versus 6A^4^ and 6B^4^-Cap^-^ (*p<*0.001; **supplementary text S3**). These results are consistent with the *rpoD* mutation increasing the probability of capsulation by generating an increase in ribosome expression.

### Manipulating components of the model generates changes in 6B^4^ capsule switching

To test whether the ribosome-RsmAE model (**fig. 6A**) underpins switching in 6B^4^, components of the model were manipulated to bias the switch in favour of ribosomes (Cap^+^) or RsmAE (Cap^-^).

The switch was tipped in favour of ribosomes by decreasing RsmAE activity in two ways. Firstly, *rsmA1* and *rsmE* were individually deleted from 6B^4^, giving 6B^4^**-Δ***rsmA1* and 6B^4^-**Δ***rsmE* (**supplementary text S2**). A capsule counting assay revealed increases in capsulation (**fig. 6B**); in particular, deletion of *rsmA1* resulted in significantly higher levels of capsulation (one-sided *t*-test *p*=0.001112), bringing the percentage of capsulated cells to almost 100%. Secondly, the model predicts RsmAE activity to be reduced by deletion of *mvaT,* which encodes a transcriptional repressor of *rsmZ* – itself a negative regulator of RsmAE – in *Pseudomonas aeruginosa* (Brencic *et al.*, 2009). Deletion of *mvaT* (*pflu4939*) from 6B^4^ or 1B^4^ results in a significant increase in capsulation (*p*<0.01; **fig. 6C**, **supplementary text S2**).

In contrast to the above, any bias of the switch machinery in favour of RsmAE is expected to inhibit translation of *pflu3655-3657* mRNA, and thus reduce capsulation. Support for this side of the model comes from the transposon mutagenesis screen (**supplementary table S1**). Firstly, three capsule-reducing insertions were obtained in the GacA/GacS two-component sensory system, which is a negative regulator of RsmAE in Γ-proteobacteria (Lapouge *et al.*, 2008) **fig. 6A**). Inactivation of these genes is expected to increase RsmAE expression and decrease capsulation. Indeed, 6B^4^-TnCre-*gacA* and 6B^4^-TnCre-*gacS* (Cre-deleted forms of the transposon mutants, see **methods** and **supplementary text S2**) showed a complete absence of capsulation (**fig. 6D**). Secondly, four transposon insertions were identified in genes involved in the production of mature tRNAs: two in *gidA/mnmG* (*pflu6129*), one in *truA* (*pflu4189*) and one in *thiI* (*pflu0349*) (reviewed in Yacoubi *et al.*, 2012). Each of these insertions resulted in a reduction in capsulation (**fig. 6D**). While not lethal, disruption of each of the three tRNA modification genes is expected to reduce translational speed (Yacoubi *et al.*, 2012; reviewed in Shepherd and Ibba, 2015), suggesting a role for efficient translation in 6B^4^ capsulation.

The ability to increase and decrease 6B^4^ capsules by manipulating components of the 1B^4^ ribosome-RsmAE circuitry (as predicted by the model) demonstrates that the same intracellular architecture underpins switching in both genotypes.

## DISCUSSION

In this work 6B^4^ has been extensively characterised. Its phenotype and genotype have been compared with those previously reported for 1B^4^ – a strain evolved in parallel to, but independently of, 6B^4^ (Beaumont *et al.*, 2009; Gallie *et al.*, 2015). 6B^4^ and 1B^4^ populations show elevated levels of CAP-based capsule expression and colony switching (**fig. 1B-E**, **2A**). The phenotype is realised by two distinct genetic routes, culminating in a mutation in either *rpoD* (Line 6) or *carB* (Line 1). Both mutations promote increased translation of mRNA encoding positive regulators of the CAP biosynthetic machinery. These regulators also activate their own transcription, forming a positive feedback loop that results in bistable capsule expression (Remigi *et al*., 2018; **fig. 6A**).

Line 1 and Line 6 were derived from a single clonal ancestor (*P. fluorescens* SBW25). This means that the genotypes of interest, 6B^4^ and 1B^4^, share an evolutionary history of many millions of years followed by a comparatively minuscule period of several weeks of independent evolution in experimental microcosms. Given the extensive shared history, it is not surprising that the same phenotype emerged in both lineages. It is surprising, however, that such different molecular pathways generate the same phenotype.

Repeated phenotypic evolution has been documented many times in both the laboratory (*e.g.*, Riehle *et al.*, 2001; Cooper *et al.*, 2003; Fong *et al.*, 2005; Ostrowski *et al.*, 2005; Bantinaki *et al.*, 2007; Meyer *et al.*, 2012; Lindsey *et al.*, 2013) and natural populations (Nachman *et al.*, 2003; Rosenblum *et al.*, 2004; Stern and Frankel, 2013; Riveron *et al.*, 2014). In many of these examples, repeated phenotype evolution is determined by changes in the same gene or molecular pathway (*e.g.*, Meyer *et al.*, 2012; Lindsey *et al.*, 2013; Riveron *et al.*, 2014). The fact that colony switching in Lines 1 and 6 arise by different molecular routes – despite extreme shared ancestry – is surprising. At first glance, *rpoD* and *carB* seem functionally unrelated and, as such, it is natural to assign them to separate functional compartments. However, as this work shows, the two genes are connected at the level of their effects on ribosomes: both mutations increase expression of ribosomal genes (**supplementary table S7**; Gallie *et al.*, 2015; Remigi *et al.*, 2018) and thus tip the balance of the switch in favour of CAP mRNA translation (**fig. 6A**).

The precise molecular mechanisms by which *rpoD* and *carB* mutations alter ribosomal gene expression remain to be elucidated. However, it is conceivable that the *rpoD* mutation directly increases transcription from one or more ribosomal genes. A point mutation in *Salmonella typhimurium rpoD* has recently been shown to increase transcription from *rpsT* (Knöppel *et al.*, 2016). In the case of the *carB* mutation, which perturbs intracellular pyrimidine pools (Gallie *et al.*, 2015), the reported influence of nucleotide triphosphate (NTP) concentrations on *rrn* promoters may play a mechanistic role (Gaal *et al.*, 1997; Schneider *et al.*, 2002; Schneider *et al.*, 2003; Murray *et al.*, 2003; Schneider and Gourse, 2003). If cellular components show a high degree of connectivity, it follows that many other factors could also affect the switch circuitry. Possible candidates include those affecting capsule expression and identified via the transposon mutagenesis screens (*e.g., hslO, sahA, ndk*; **supplementary table 1**; Gallie *et al*., 2015).

In stark contrast to the disparate molecular evolution of 6B^4^ and 1B^4^, repeated bouts of evolution from the same immediate ancestor of the 6B^4^ switching genotype, namely, 6A^4^, resulted in re-evolution of the switching genotype by mutations solely in *rpoD* (**fig. 5A**). Similar repeated bouts of evolution from 1A^4^ (the immediate ancestor of the Line 1 switching genotype) resulted in switching types with mutations in genes encoding the determinants of pyrimidine biosynthesis (five in *carB*, one in *pyrH*; Gallie *et al*., 2015). In other words, the comparatively tiny portion of evolutionary history for which Line 6 and Line 1 diverged – several weeks compared with millions of years of common history – has a significant impact on molecular evolution.

The distinct classes of switcher mutations in Lines 1 and 6 are likely to result from positive epistatic interactions: while both types of switch-causing mutations presumably arise in both backgrounds, *rpoD* mutations provide a significant growth advantage in 6A^4^ (and, by extension, *carB/pyrH* mutations in 1A^4^). The growth advantage afforded by *rpoD* mutations in 6A^4^ can be seen in **fig. 5D**, where each of three *rpoD* mutants grows more quickly and to a higher final density than ancestral 6A^4^ in shaken KB. The advantage of the t1682c *rpoD* mutation (that from 6B^4^) disappears in the absence of the previous six mutations (SBW25 grows faster than SBW25-*rpoD\** (**fig. 5D**)). Precisely which of the first six mutations in the Line 6 evolutionary series contribute to the observed epistatic effect remains to be tested. However, the two *nlpD* mutations immediately preceding the *rpoD* mutation are prime candidates for two reasons. Firstly, *nlpD* is the only locus that is mutated in Line 6 but not Line 1 (**fig. 1B**, **2A**). Secondly, *nlpD* is immediately upstream of *rpoS* (*pflu1302*), which encodes the stationary phase sigma factor RpoS (Σ^38^). RpoS and RpoD (together with other sigma factors) compete for binding of core RNA polymerase (Ishihama, 2000; Mauri and Klumpp, 2014), and so their relative intracellular concentration affects the expression level of their respective regulons (Gross *et al.*, 1998; Mauri and Klumpp, 2014). It is possible that the *nlpD* mutations, in addition to altering colony morphology *via* a reduction of NlpD/AmiC activity, also alter the expression of *rpoS*. Indeed, a promoter for *rpoS* has previously been reported within *E. coli* and *P. aeruginosa nlpD* (Takayanagi *et al.*, 1994; Lange and Hengge-Aronis, 1994; Kojic and Venturi, 2001). A change in RpoS concentration could conceivably set the stage for compensatory mutations in RpoD.

Understanding the molecular bases of adaptive phenotypes continues to present significant challenges even when aided by high-throughput genomic technologies. As shown here and elsewhere (Larsen *et al.*, 2008; Gallie *et al.*, 2015; Bershtein *et al.*, 2015; Grenga *et al.*, 2017; Carvalho *et al.*, 2018), mutations – particularly those in central metabolism – can have complex effects that extend well beyond the immediate neighbourhood of gene function. That point mutations in two seemingly unrelated genes (*rpoD* and *carB*) can generate stochastic capsule switching draws attention to the interconnectedness of cell physiology and highlights the extensive mutational opportunities available to evolution.

## MATERIALS AND METHODS

### Bacterial strains, plasmids and media

Details of bacterial strains and plasmids used are provided in **supplementary text S2**. Unless otherwise stated, *P. fluorescens* strains were grown for 24 hours at 28^°^C in shaken 30 mL glass microcosms containing 6 mL King’s Medium B (KB; Ward *et al.*, 1954). Where stated, uracil L-arginine hydrochloride and/or guanine hydrochloride (Sigma-Aldrich) were added to the medium. Cells were plated on KB or Lysogeny Broth (LB) containing 1.5% agar. Antibiotics were used at the following concentrations: tetracycline (12.5 μg mL^-1^; Tc); kanamycin (100 μg mL^-1^; Km); nitrofurantoin (100 μg mL^-1^; NF).

### Microscopy

Cell microscopy was performed using a Zeiss Axiostar Plus bright field microscope. A dissection microscope was used for colony images. Microscopy images were cropped and processed in Preview or Microsoft Word as indicated in figure legends.

### Capsule counting assay

Capsule staining and the counting assay were performed as previously described (Gallie *et al.*, 2015). Briefly, for each strain to be assayed, three (for the Cre-deleted transposon mutant assay in **fig. 6D**) or five (all others) single colonies were grown to stationary phase in KB cultures. Cultures were transferred to fresh KB and grown to mid-exponential phase. Cells from each culture were stained with 1:10 diluted India ink (Pébéo) and photographed under bright field x60 magnification. Capsule expression was recorded manually for 500 cells *per* replicate (≤100 cells assayed *per* photograph). Average proportions of Cap^+^ cells were determined and statistical analyses performed.

### Gene deletions and mutation construction

Gene deletions were constructed in the SBW25 background by pUIC3-mediated two step allelic exchange as described elsewhere (Zhang and Rainey, 2007). For further details of genetic constructs see **supplementary text S2**.

### Construction and analysis of *wcaJ-lacZ* transcriptional fusions

Wild type *wcaJ* was PCR-amplified and ligated into pUIC3 (Rainey, 1999) immediately upstream of promoter-less *lacZY* (see **supplementary text S2**). The construct was used to transform *E. coli* DH5α-Λ*pir* and transferred to 6A^4^ and 6B^4^ *via* tri-parental conjugation (with a helper strain carrying pRK2013). Successful transconjugants were purified, giving 6A^4^-*wcaJ-lacZ* and 6B^4^-*wcaJ-lacZ*. Single colonies of these constructed genotypes were grown at 26^°^C for 48 hours on LB+Tc+X-gal (60 μg mL^-1^) plates prior to microscopic analysis.

### Transposon mutagenesis

6B^4^ was subjected to random mutagenesis as described in Giddens *et al.*, 2007. Approximately 10,000 transposon mutants from eleven independent conjugations were screened on LB+Km, on which 6B^4^ mutants typically form opaque colonies after ∼72 hours. Mutants that formed translucent or otherwise different colonies were selected and screened microscopically for obvious alterations in capsule expression. Mutants of interest were then purified and the insertion site determined by AP-PCR. In selected strains, the bulk of the transposon was deleted leaving 189 base pair at the insertion site (“TnΔ-” genotypes) and eliminating polar effects (Giddens *et al.*, 2007).

### Isolation and analysis of extracellular polysaccharide (EPS)

Extracellular polysaccharide (EPS) was isolated from SBW25, 6A^4^ and 6B^4^. Each genotype was grown on KB agar (28^°^C, 48-96 hours). Cellular material was resuspended in 12 mL of 1 M NaCl to give an OD_600_ of ∼3.5, vortexed for 40 min and centrifuged (30 min, 4168 *g*). EPS was precipitated from supernatants by addition of 3 volumes isopropanol. Following re-suspension of pelleted EPS (40 min, 4168 *g*) in 0.5 mM CaCl_2_, RNase (0.1 mg mL^-1^) and DNase (1.2 units mL^-1^) were added and samples incubated at 37^°^C overnight. The next day samples were supplemented with citrate buffer (pH 4.8, 50 mM) and, to eliminate any cellulose, treated with cellulase (ICN Biomedicals Inc.; 0.15 mg mL^-1^, 2 hours at 50^°^C). After addition of 0.5 mg mL^-1^ Proteinase K, EPS was precipitated by addition of 3 volumes isopropanol and pelleted (40 min, 4168 *g*). EPS pellets were dissolved in dH_2_O (50^°^C, overnight). Finally, samples were dialyzed against dH_2_O for 48 hours (SnakeSkin^®^ dialysis tubing, Thermo Scientific). The Callaghan Research Institute (New Zealand) performed EPS analysis. Samples were freeze-dried and weighed, and a colorimetric total sugar assay performed in duplicate for each EPS isolation.

### Genome Sequencing of 6B^4^

A 6B^4^ colony was grown overnight in KB. Cap^+^ and Cap^-^ fractions were separated by centrifugation and genomic DNA isolated from each fraction using the CTAB method. Equal quantities of each isolated DNA were mixed, and whole genome re-sequencing was performed (Illumina; Massey University, NZ). Point mutations were identified by aligning 36 base pair sequence reads to the SBW25 genome (Silby *et al.*, 2009) *via* SOAP (short oligonucleotide alignment program (Li *et al.*, 2008)) and ELAND (Illumina, Inc.). Insertions and deletions were identified by analysing genomic regions with unusual coverage and BLAST analysis of discarded sequences. Genome sequence files available on request.

### Re-evolution of switchers from 6A^4^

Nine independent switcher genotypes were isolated from 6A^4^ according to the REE protocol (Beaumont *et al.*, 2009). Each switcher was purification streaked and *rpoD* sequenced.

### Growth curves and analysis

Single colonies were grown for each strain of interest (KB agar, 26^°^C, 48 h). Eight colonies *per* strain were used to inoculate 200 μL KB in wells of a 96-well plate. The plate cultures were grown for 24 hours at 26^°^C, 200 rpm. Each well was then mixed by pipetting and 2 μL culture transferred to 198 μL fresh KB. This second plate was then incubated at 26^°^C for 72 hours in a BioTek Epoch 2 plate reader, and the OD_600_ of each well measured at five minute intervals (5 seconds of 3 mm orbital shaking preceding each read). Data from each well was plotted, and the mean and standard error of eight wells *per* strain (minus absorbance in the media control wells) used to draw **fig. 5D**. V_max_ (maximum growth rate) and lag time were calculated using a sliding window of six time points during exponential growth (between 1-24 hours, based on observation of growth curves) using Gen5 Software version 3.00.19.

### RNA-seq analysis

For each of 6A^4^ and 6B^4^, three single colonies were grown overnight in independent KB microcosms, diluted 1:1000 into 20 mL KB in 250 mL flasks and grown to mid-exponential phase (∼OD_600_ of 0.4-0.6). Total RNA was then harvested from each culture; for 6A^4^ cultures, 100 μL culture were mixed with 900 μL KB, pelleted and resuspended in 1 mL of RNAlater^®^ (Ambion^®^). For 6B^4^ cultures, 100 μL was separated into Cap^+^ and Cap^-^ fractions by centrifugation, and each was resuspended in 1 mL RNAlater^®^. All mRNA extractions proceeded using a RiboPure^TM^ Bacteria Kit (Ambion^®^). Normalized mRNA-seq library preparation, followed by 100 base pair paired-end Illumina HiSeq 2500 sequencing, was performed by the Australian Genome Research Facility (Brisbane, Australia; GEO submission number GSE116490). The data was analysed with Bowtie2 (Langmead and Salzberg, 2012), HTSeq (Anders *et al.*, 2015) and the R-package DESeq2 (Love *et al.*, 2014). First, RNA-seq datasets were mapped to the SBW25 genome (downloaded from GenBank under the accession number NC_012660) via Bowtie2 with default settings. The coverage *per* gene of the genome mapping was determined with HTSeq. The gene annotation for HTSeq was also downloaded from the SBW25 GenBank entry. Then, differentially expressed genes were identified by applying DESeq2. The standard workflow inhttps://bioconductor.org/packages/release/bioc/manuals/DESeq2/man/DESeq2.pdf was used, except that the alpha parameter was set to 0.3 to reduce the number of genes that were falsely classified as not significantly differentially expressed between the different morphotypes. Three comparisons were made: 6A^4^ versus 6B^4^-Cap^-^, 6A^4^ versus 6B^4^-Cap^+^, and 6B^4^-Cap^-^ versus 6B^4^-Cap^+^ (**supplementary tables S3-S5**). The corresponding comparisons for Line 1 are available elsewhere (Gallie *et al.*, 2015).

### Statistical analyses

To detect differences in capsulation levels or nucleotide concentrations between two strains, two-sample *t*-tests (parametric or Welch) or, where normality assumptions were violated, Wilcoxon rank sum tests were applied. To detect differences in capsulation levels across three strains (while testing over-expression genotypes), one-way ANOVA or, where normality assumptions were violated, Kruskal Wallis tests were used. Exact binomial tests were used to detect differences in ribosomal gene expression between morphotypes in the RNA-seq data (see also **supplementary text S3**). All analyses were performed in R version 3.3.3. On graphs: * =0.05<*p<*0.01, * * =0.01<*p<*0.001, * * * =*p*<0.001.

## ACKNOWLEDGMENTS AND FUNDING INFORMATION

The raw RNA-seq data is deposited at GEO (submission number GSE116490). This work was supported by The Marsden Fund (all authors) and the Max Planck Society (JG, FB, PBR).

## AUTHOR CONTRIBUTIONS

JG and PBR designed the study; JG, PR, GCF and SN performed the experiments; FB and JG analysed the data; JG wrote the manuscript with contributions from all authors.

